## Supplementary Data Index File for "Transcriptome-wide analysis of the function of Ded1 in translation preinitiation complex assembly in a reconstituted in vitro system"

**Supplementary file 1**

**Excel file containing results and analyses from Rec-Seq experiments**

**Spreadsheet 1**, 'mRPF Ded1 for Fig2', **column 2-4,** for means of normalized mRPFs of -Ded1 (mRPF_0nM_Ded1), 100 nM Ded1 (mRPF_100nM_Ded1), 500nM Ded1 (mRPF_500nM_Ded1), **column 5-7,** for log2 values of ΔmRPF100/0nM Ded1 (FC_100/0nM_Ded1), ΔmRPF500/0nM (FC_500/0nM_Ded1) and ΔmRPF500/100nM (FC_500/100nM_Ded1), **column 8-10,** FDR 100/0nM Ded1 (FDR_100/0nM_Ded1), FDR 500/0nM Ded1 (FDR_500/0nM_Ded1), FDR 500/100nM Ded1 (FDR_500/100nM_Ded1), for 3052 mRNAs described in Fig. 2A-C.

**Spreadsheet 2,** 'mRNA features for Fig3-5', For mRNA features including CDS length, 5'UTR length, context scores described in (Martin-Marcos et al. 2017) and PARS score analysis of 5' UTRs and 5' end of CDS of 2679 genes, conducted as described in (Sen et al. 2015) (NA for 5'UTR length, context score or PARS scores not annotated in original research).

**Spreadsheet 3,** "RiboSeq *ded1-cs* for Fig4', Reanalysis of ribosome profiling results of (Sen et al. 2015), **column 2-10,** for log2 values of WT mRNA (mrnaWT), WT RPF (riboWT), *ded1-cs* mRNA (mrnaded1_cs), *ded1-cs* RPF (riboded1_cs), ΔmRNA *ded1-cs* (mrnaChange), ΔRPF *ded1-cs* (riboChange), WT TE (teWT), *ded1-cs* TE (teded1-cs) and ΔTE *ded1-cs* (teChange), **column 11,** log10 FDR for comparing WT and ded1-cs TEs ( logFDR), **column 12-13,** for gene names and description.

**Spreadsheet 4,** 'iRPF_Ded1_for_FIg5', **column 2-4,** means of normalized iRPFs of -Ded1 (iRPF_0nM_Ded1), 100 nM Ded1 (iRPF_100nM_Ded1), 500nM Ded1 (iRPF_500nM_Ded1), **column 5-6,** for log2 values of ΔiRPF100/0nM Ded1 (FC_100/0nM_Ded1), ΔiRPF500/0nM (FC_500/0nM_Ded1), **column 7-8,** FDR 100/0nM Ded1 (FDR_100/0nM_Ded1), FDR 500/0nM Ded1 (FDR_500/0nM_Ded1) for 3052 mRNAs described in Fig. 5I-J.

**Spreadsheet 5,** 'uRPF_Ded1_for_Fig6, **column 2-4,** means of normalized uRPFs of -Ded1 (uRPF_0nM_Ded1), 100 nM Ded1 (uRPF_100nM_Ded1), 500nM Ded1 (uRPF_500nM_Ded1), **column 5-6,** for log2 values of ΔuRPF100/0nM Ded1 (FC_100/0nM_Ded1), ΔuRPF500/0nM (FC_500/0nM_Ded1), **column 7-8,** FDR 100/0nM Ded1 (FDR_100/0nM_Ded1), FDR 500/0nM Ded1 (FDR_500/0nM_Ded1) for 3052 mRNAs described in Fig. 6B-C.

**Spreadsheet 6,** 'mRPF_eIF4A_for_Fig7, **column 2-3,** mean of normalized mRPFs of 5000nM (mRPF_5000nM), 500nM eIF4A (mRPF_500nM), **column 4,** for log2 values of ΔmRPF 5000/500 nM eIF4A (FC_5000/500nM), **column 5,** FDR 5000/500 nM eIF4A for mRNAs described in Fig.7A.

**Spreadsheet 7,** 'Input_mRNA' for the RNA-Seq reads (mRNA) and mRNA-density calculated as reads per nucleotide (mRNA_density).

Martin-Marcos P, Zhou F, Karunasiri C, Zhang F, Dong J, Nanda J, Kulkarni SD, Sen ND, Tamame M, Zeschnigk M et al. 2017. eIF1A residues implicated in cancer stabilize translation preinitiation complexes and favor suboptimal initiation sites in yeast. *Elife* **6**.

Sen ND, Zhou F, Ingolia NT, Hinnebusch AG. 2015. Genome-wide analysis of translational efficiency reveals distinct but overlapping functions of yeast DEAD-box RNA helicases Ded1 and eIF4A. *Genome Res* **25**: 1196-1205.
